# Variant annotation across homologous proteins (“Paralogue Annotation”) identifies disease-causing missense variants with high precision, and is widely applicable across protein families

**DOI:** 10.1101/2023.08.07.552236

**Authors:** Nicholas Li, Xiaolei Zhang, Erica Mazaika, Pantazis Theotokis, Mikyung Jang, Mian Ahmad, George Powell, Henrike O. Heyne, Dennis Lal, Paul JR Barton, Roddy Walsh, Nicola Whiffin, James S Ware

**Author notes:** Corresponding author James Ware, National Heart & Lung Institute & MRC Laboratory of Medical Sciences, Imperial College London, London W12 0HS, UK. these authors contributed equally (Nicholas Li, Xiaolei Zhang).

## Abstract

**Background:** Distinguishing pathogenic variants from those that are rare but benign remains a key challenge in clinical genetics, especially for variants not previously observed and characterised in humans. *In vitro* and *in vivo* functional characterisation are typically resource intensive, and model systems may not accurately predict influence on human disease. Many *in silico* tools have been developed to predict which variants are disease-causing, but typically lack precision. Here we demonstrate the applicability of a framework, called Paralogue Annotation, that draws on information from previously-characterised variants in homologous proteins to predict whether variants in a gene of interest are likely disease causing.

**Methods:** We assessed the performance of Paralogue Annotation through three orthogonal approaches: (1) comparison to established *in silico* variant prediction tools using 47,016 missense variants from ClinVar across 3,524 genes representing a broad range of diverse protein classes, by calculating precision and sensitivity; (2) evaluation against large-scale functional assays of variant effect; and (3) comparing odd ratios calculated from case-control association tests for inherited cardiac arrhythmia syndromes, and neurodevelopmental disorders with epilepsy, stratifying variants by Paralogue Annotation.

**Results:** Paralogue Annotation correctly annotates 4,328 ClinVar pathogenic variants, with 245 false positives, yielding a precision of 0.95. This increases to 0.99 with more stringent annotation parameters (requiring greater conservation of amino acids between annotated orthologues) at the expense of sensitivity. Compared to established tools, Paralogue Annotation has higher precision for the identification of pathogenic variants, albeit with lower sensitivity across diverse test sets. Extending the technique by transferring annotations between homologous protein domains, rather than full-length protein paralogues, increases sensitivity. Rare variants predicted pathogenic by Paralogue Annotation were more strongly disease-associated (increased odds ratio) than unstratified rare variants for six out of eight genes tested with case-control cohort approaches.

**Conclusions:** We demonstrate that Paralogue Annotation has high precision for predicting pathogenic missense variants, providing a useful line of evidence for clinical variant interpretation that is complementary to other approaches in use. As the number of characterised variants increases in reference datasets such as ClinVar, Paralogue Annotation will further increase in sensitivity and applicability.

## Background

Distinguishing variants in the human genome that have the potential to cause monogenic disease remains a key challenge in genomic medicine. The American College of Medical Genetics and Genomics and Association for Molecular Pathology (ACMG/AMP) provides a framework for variant interpretation^1^ that has been widely adopted internationally, in which different lines of evidence for or against pathogenicity are combined in a semi-quantitative decision matrix. This matrix is used to estimate the likelihood that a variant is disease-causing, given that it has been observed in an individual with disease. Lines of evidence include direct observations of the variant of interest in humans in health or disease, including assessment of the concordance with known inheritance patterns and mechanisms of disease, and studies of the functional effects of the variant *in silico*, and in *in vitro* or *in vivo* model systems.

When a variant has not been observed in humans, nor functionally characterised, the framework considers evidence drawn by inference from other variants that might be considered functionally equivalent. For example, if loss-of-function is established as the mechanism of disease, then any predicted null variant that is sufficiently rare will be considered likely pathogenic, in the absence of specific contradictory data^2^ (PVS1, PM2). Similarly, if a missense variant that has not been previously described lies in a gene or region in which neutral missense variants are rarely observed (PP2, PM1), or affects the same amino acid residue as a known pathogenic variant (PS1/PM5), then this is evidence favouring pathogenicity.

Although not yet formalised in the overarching ACMG/AMP framework, there is another context in which substitutions may be considered as functionally equivalent: when they impact the equivalent position of different members of a protein family (paralogues). For example, if a substitution in an ion channel expressed in the brain causes an inherited epilepsy syndrome, then one might hypothesise that an equivalent substitution in a paralogous protein expressed in the heart would cause an inherited arrhythmia syndrome, through an equivalent molecular mechanism in another excitable tissue. We have previously validated this approach using genes associated with inherited cardiac conditions and their paralogues, and found that such paralogue annotation (PA) has a very high positive predictive value for pathogenicity ^3,4^, and others have subsequently independently confirmed these findings.^5,6,7,8,9^

Some gene- and disease-specific adaptations of the ACMG/AMP framework have since instantiated equivalent rules for specific closely-related proteins. For example, ClinGen’s RASopathy Expert Panel advises that PS1, PM1 and PM5 can be applied for “analogous residue positions/regions” across specific groups of proteins such as BRAF/RAF1 and HRAS/KRAS/NRAS ^10^.

There is a continuum of relatedness amongst proteins. Our previous work focused on transferring annotations between evolutionarily closely-related paralogues, but the same principle might be applied to more distantly related proteins that have diverged in function, but share common domains where equivalence is preserved. For example, many of the distinct families of paralogous ion channels that we studied previously can be grouped into a cation channel superfamily, comprising proteins with diverse ion selectivity and gating mechanisms, but shared features including a tetrameric structure and homology of the pore-forming regions PF00520^11^.

Here we evaluate the broad applicability of Paralogue Annotation. We characterise for the first time the precision of annotations made between direct paralogues exome-wide, by assessing performance both against gold-standard reference sets of interpreted variants, and also using additional approaches that are independent of a gold-standard benchmark classification. We then extend the approach to consider annotations across homologous protein domains that are shared by more distantly-related proteins. We demonstrate this approach is widely applicable and highly informative, and provides a useful line of evidence to enhance the interpretation of previously unseen genetic variants.

## Methods

### Paralogue Annotation procedure and implementation

A schematic of the PA approach is shown in **Fig. 1 (A)**. Given a query variant (or set of variants) in a gene of interest, we first identify paralogues of that gene. A multiple sequence alignment (MSA) of members of the protein family identifies the residues in each paralogue that are homologous to the position(s) of the query variant(s). Here, we used predefined paralogues and MSAs from Ensembl Compara^12^. We then search for any known (human) variants at these equivalent aligned positions. For this, we used ClinVar^13^ (version 20190114) as the most comprehensive variant catalogue. If variation at these proteins is known to be pathogenic, we transfer the consequence annotation across the alignment between paralogues, inferring that an equivalent substitution at the query variant may also be pathogenic.

**Fig. 1.**
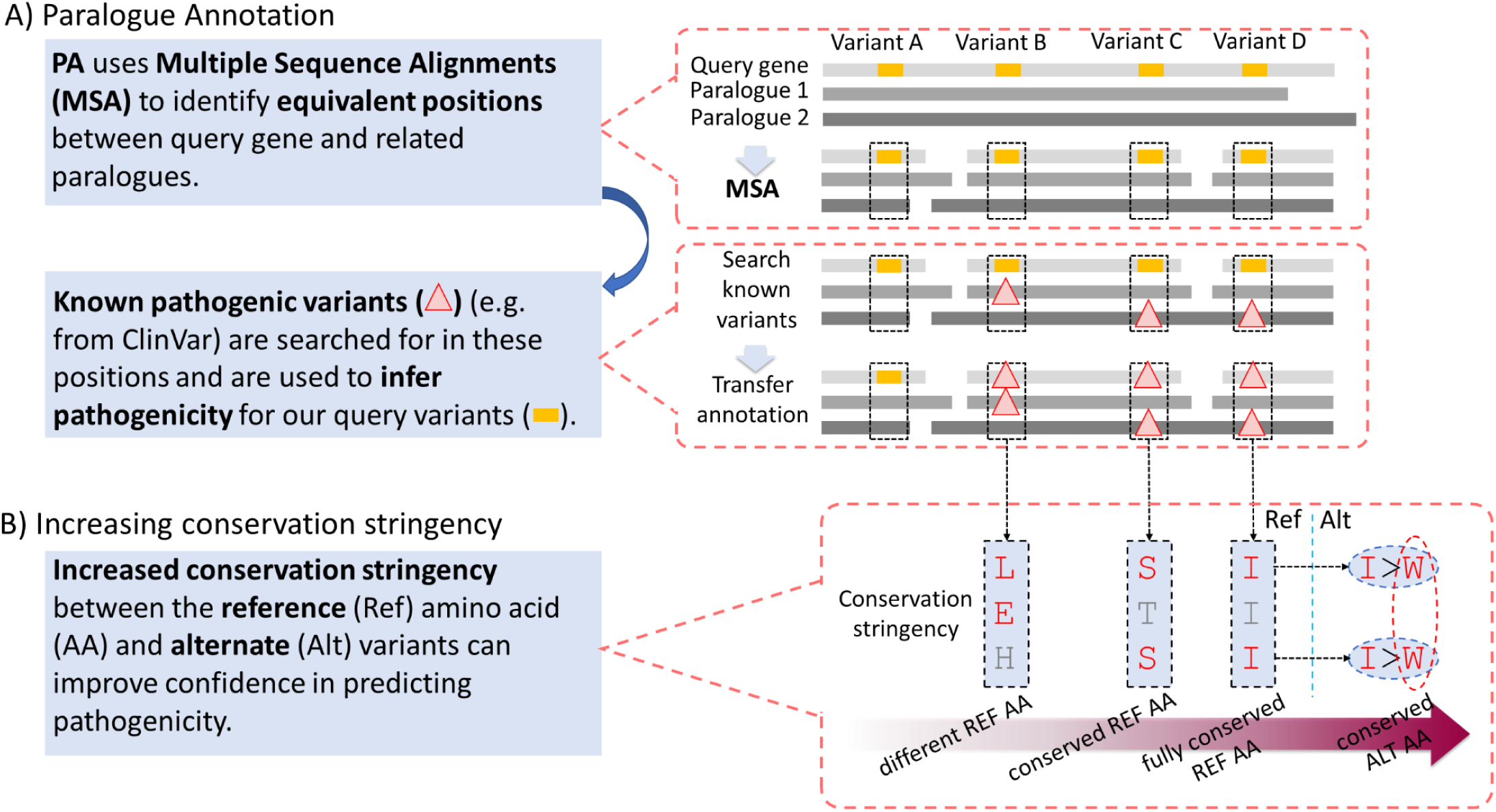
Paralogue Annotation overview. (A) Paralogues of the query gene of interest are aligned to identify equivalent positions (dotted boxes) to the variant of interest (yellow box). Any known disease-causing variants (red triangles) in those equivalent positions are identified. Here Variant A does not have any aligned disease-causing variants while Variants B, C and D do. Their positions are mapped back to the query gene and their annotations transferred, inferring that variants at these sites are likely to also be disease-causing. (B) The conservation between the reference (ref) and alternate (alt) amino acids (AA) in the query gene and paralogues can be used to filter annotations to improve confidence in calling pathogenicity. With no conservation stringency, all alignments will be used regardless of AA similarity (Variant B). By increasing the stringency, only alignments where there are conserved ref AA between the query gene and the paralogue with the annotation are used (Variant C). Increasing the stringency further, full family-wide ref AA conservation is required before annotation transfer (Variant D). In addition, conservation between the alt AA in the query gene and the paralogue can also be considered.

A Perl (Version 5.10.1) plugin for Ensembl’s Variant Effect Predictor (VEP) ^14^ was developed to allow flexible use of PA on query variants listed in a Variant Call Format (VCF) file. The plugin utilises pre-aligned paralogues from Ensembl’s Compara database ^15^ to return any equivalent amino acids (AA) and their coordinates. A reference set of known disease-causing variants is then searched for variants at these positions. In this study, ClinVar variants with a clinical significance of Pathogenic/Likely Pathogenic (P/LP) were used, but alternative sets of variants with well-characterised links to human disease can be used instead or as well. We note that the pre-defined paralogue alignments by Ensembl sometimes caused particular issues when transferring annotations across equivalent positions, such as overlapping genes or read-through transcripts that were defined as self-paralogues (**Additional File 1**).

### Characterising protein families of disease-associated and non-disease-associated genes in the human genome

To determine the relevance of PA to human disease genes, we counted paralogues of genes with and without links to human disease. This was done using human genome build GRCh38 data analyzed from Ensembl Biomart (Ensembl version 93: Jul2018) in R with the BiomaRt package from Bioconductor^16,17^. All protein coding genes were retrieved. Disease-associated genes were here defined as those that contained at least one Pathogenic or Likely Pathogenic (P/LP) variant in ClinVar. A two-sample Kolmogorov–Smirnov test was used to compare distributions.

### ClinVar reference data

We downloaded 440,203 variants in 5,863 protein coding genes from ClinVar version 20190114, build GRCh37^18^ from ClinVar’s FTP site (ftp.ncbi.nlm.nih.gov/pub/ClinVar). We used 30,334 missense variants classified as Pathogenic or Likely Pathogenic and 16,682 missense variants classified as Benign or Likely Benign with no conflicting interpretations to analyse the performance of PA.

### Utilizing conservation stringency to increase confidence in functional equivalence of variants at homologous positions

For each query variant, the reference amino acid conservation across the protein family alignment can be considered via an order of increasingly stringent filters (**Fig. 1b**) to increase confidence that homologous residues are structurally and functionally equivalent. At the lowest stringency conservation is not evaluated, and we examine all aligned positions (“*different ref AA”* in **Fig. 1b**). At the next level of stringency we examine only alignments where the reference amino acids are conserved between the query protein and the source protein, but not requiring conservation of the ref AA across the whole paralogue family(“*conserved ref AA”*). Thirdly we examine only alignments where the reference AA is conserved across all members of the protein family (“*fully conserved ref AA”*). Finally, in addition to considering conservation of the reference amino acids, we can also consider whether the substituting (alternate) amino acids are equivalent; here we require that the amino acid substitutions are identical in both query protein and source protein in order to transfer annotations (“*conserved alt AA”*). This step could be applied independently following any of the aforementioned steps, but in **Fig. 1b** it is applied alongside filters requiring conservation of the reference AA as an example. We further evaluated Para Z scores, as described by Lal *et al.* ^19^, as a more granular quantification of paralogue conservation across families.

All of these additional conservation stringencies are not needed for PA to function but are supplemental in improving its performance.

### Comparing the performance of Paralogue Annotation to other computational tools

When a variant is coincident with a known pathogenic variant in a paralogue we hypothesise that this is evidence for pathogenicity, but we do not expect that the absence of a previously characterised variant in a paralogue would constitute evidence against pathogenicity given that phenotypic consequences remain uncharacterised for the majority of possible missense variants in the human genome. Therefore, Paralogue Annotation is not conceptually equivalent to tools that attempt to classify variants as either pathogenic or benign.

We define variants as either “test positive” or “test negative” according to the presence/absence of paralogue annotation (i.e. when a query variant aligns with a P/LP reference variant in one or more paralogues). To analyse performance we focus on measures related to a positive test result, where interpretation is most meaningful and relevant: 1) The Positive Predictive Value (PPV; also known as precision) measures the proportion of variants correctly predicted as being pathogenic out of all variants that were predicted positive; and 2) the sensitivity (also known as recall, or the true positive rate) measures the proportion of actual pathogenic variants that are correctly predicted as such.

A true positive (TP) indicates that a ClinVar P/LP query variant is annotated with a homologous P/LP variant, and a false positive (FP) indicates that a ClinVar B/LB variant is annotated with a homologous P/LP variant.

The Precision and Sensitivity are therefore calculated as follows:

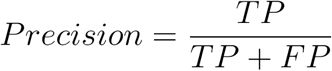

and

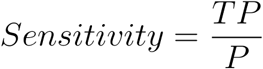

where *P* is the total number of P/LP query variants used in the analysis.

P-values for statistical significance are calculated via a Fisher’s exact test on a 2x2 contingency table. All calculations were performed in R (version 3.6.3)

The performance of PA as a classifier was compared to AlphaMissense^20,21^), EVE^22^, REVEL ^23^, SIFT^24^, M-CAP ^25^ and CADD ^25^. Only variants that were able to be annotated by all methods were used during comparisons.

### Functionally characterised variants from large-multiplex assays

As an additional test set, we used datasets with multiplex assays of variant effect (MAVEs) curated from ProteinGym^26^. In total, 64 genes available with both MAVE assays and the baseline paralogue annotation (without conservation filters) are included. For some genes that have more than one assay available, we chose the one with the lowest average fitness scores, assuming it’s the scenario where mutations can induce the worst impact on fitness.

### Assessing performance using case-control cohort data

As an orthogonal approach that is not reliant on the accuracy of pathogenicity assertions in the ClinVar “gold-standard” reference data, nor reliant on the validity of functional assays as surrogates for *in vivo* pathogenicity, we used cohort approaches to directly assess the strength of association between rare variants and disease for (i) all rare variants, (ii) rare variants that occurred at paralogous conserved sites (identified using a positive Para Z score^27^), (iii) rare variants that were also predicted as pathogenic by PA, (vi) rare variants predicted as pathogenic by PA with a conserved Ref AA and (v) rare variants predicted as pathogenic by PA with a fully conserved Ref AA.

Case/Control Odds Ratios (OR) were calculated for each group of variants, using cohorts diagnosed with LQTS ^28^, Brugada Syndrome (BrS) ^29^, or Neurodevelopmental Disorder with Epilepsy (NDD+E) ^30^ , and considering gnomAD v2.1.1^31^ as a control population. Rare variants in LQTS and BrS were defined as any that had a gnomAD Minor Allele Frequency (MAF) of ≤ 8.2 × 10^-6^ and 1.0 × 10^-5^ respectively, as previously described^32^. Rare variants in NDD+E were defined as those not present in DiscovEHR^33^. For each comparison the same definition was applied to Gnomad controls.

OR were calculated from the cases and controls for all rare variants and compared to OR calculated from rare variants that were also predicted as pathogenic by PA.

OR were calculated by:

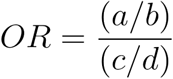

where *a* is the number of cases with variants that are rare (and predicted to be pathogenic), *b* is the number of controls with variants that are sufficiently rare (and predicted to be pathogenic), *c* is the number of cases without variants that are sufficiently rare (and predicted to be pathogenic), and *d* is the number of controls without variants that are sufficiently rare (and predicted to be pathogenic).

The 95% confidence intervals for OR values were calculated according to Altman ^34^ via:

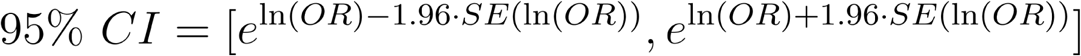

where:

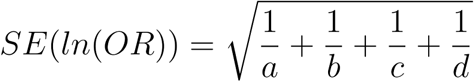

Finally, we interrogated an additional dataset of *de novo* mutations (DNM) identified by trio sequencing of individuals with autism spectrum disorder (ASD)^35^ and their parents, alongside control trios. Here we compared the DNM rate exome-wide in individuals with ASD vs controls for all DNMS, and for DNMs annotated as likely pathogenic by PA. Rates were compared using a two sampled poisson hypothesis test.

### Annotating homologous positions across Pfam domains

To improve the sensitivity of this framework, we evaluated an extension of the method that considers a broader definition of homology to identify a potentially much larger pool of “equivalent positions’’ in evolutionarily related proteins. Instead of looking at only the immediate protein family (paralogues), we consider Pfam domains, since specific functional domains can be identified as homologous between much more distantly-related proteins. The procedure described above was repeated, but substituting Compara protein-family alignments with Pfam metadomain alignments^36^. Since these alignments are not represented in Compara, custom Python and R scripts were used, rather than a VEP plugin, to annotate test variants using the ClinVar reference data as before.

## Results

### Human disease genes often have paralogues that can inform variant interpretation

Most human disease genes, as defined by the presence of at least one known (likely) pathogenic variant in ClinVar, have one or more paralogues, and could potentially be annotated using this approach. 72% (14,514/20,158) of all Ensembl protein coding genes have at least one paralogue (**Fig 2a**), with a mean of 4.5 paralogues per gene, and the largest family comprising 49 paralogous Zinc Finger proteins. Amongst 5,117 genes associated with human disease in ClinVar the proportion with one or more paralogue is similar (71%, 3,621/5,117), with slightly fewer paralogues per gene (mean 4.1) as expected. To date, about half of these (54%, 1,947/3,621) have at least one paralogue that has also been implicated in monogenic human disease and may therefore be informative for clinical variant interpretation using the proposed approach. This proportion is anticipated to increase with ongoing discovery of disease-associated genes and disease-causing variants.

**Fig. 2.**
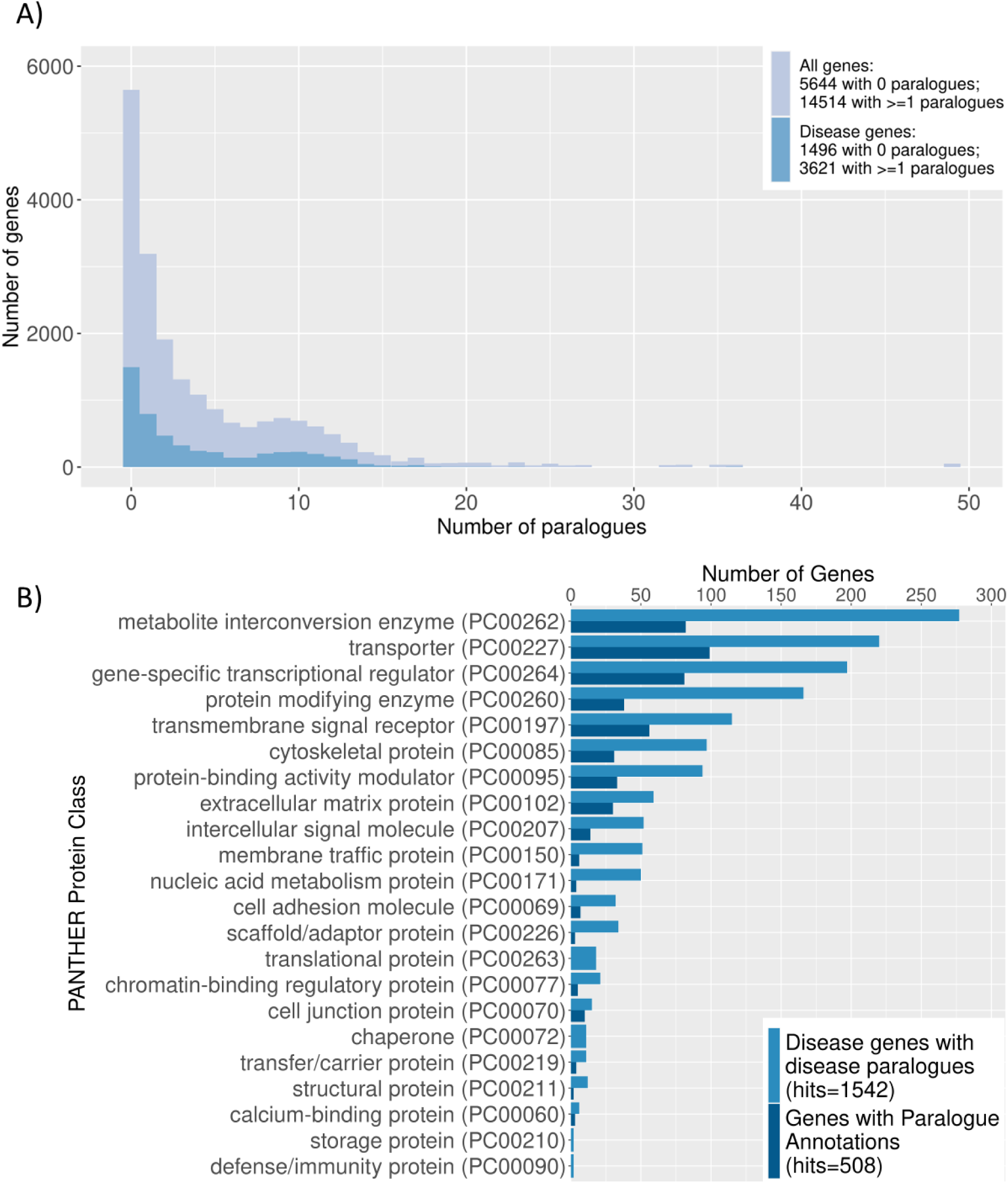
Summary of human paralogues in the human genome. *a)* Most human protein-coding genes, including those known to be associated with disease, have at least one paralogue. A histogram shows the distribution of the number of paralogues for all human protein-coding genes (light blue), and for genes known to be associated with monogenic disease (dark blue; defined as genes with at least one pathogenic or likely pathogenic variant in ClinVar). 71% of disease-associated genes have at least one paralogue, which may be informative for paralogue annotation. ***b)* Paralogue Annotation is applicable to genes encoding a diverse range of protein classes.** The distribution of protein classes encoded by these genes is shown (defined using the PANTHER classification): in light blue - ClinVar disease genes that have paralogues which are also involved in disease; in dark blue - genes that had at least one variant predicted to be pathogenic by Paralogue Annotation.

Disease-associated genes with disease-associated paralogues span a wide range of protein classes, assessed using PANTHER classifications (**Fig. 2b light blue bars**, for list of genes see **Additional File 2**). This suggests that this approach is widely applicable across diseases with diverse underlying pathological mechanisms, and is not restricted to closely related ion channels that have been evaluated previously^3,4^. PANTHER assigns classifications for 1,542 out of 1,947 disease genes with disease-associated paralogues, which span 22 protein classes. The most highly represented protein classes were metabolite interconversion enzymes (277 hits), which includes enzymes such as hydrolases, isomerases, and ligases, followed by families of transporters (220) and gene-specific transcriptional regulators (197). These latter classes encompass channel proteins and transcription factors respectively. The classes with the fewest genes with paralogues were calcium-binding (6), storage (2) and defense/immunity proteins (2).

While 1,947 disease genes have disease-associated paralogues, and could therefore theoretically be assessed using PA, some do not have missense P/LP variants in alignable regions in ClinVar to date. We therefore repeated these analyses for the subset of genes with variants predicted pathogenic by PA, with similar results (shown as **dark blue bars** in **Fig. 2b**).

### Paralogue annotation has high precision for detection of pathogenic variants in ClinVar

When PA is applied to the 30,334 P/LP ClinVar variants, 4,328 are correctly predicted to be pathogenic (14%). Conversely, only 245 B/LB variants are incorrectly predicted to be pathogenic out of 16,724 (1.4%). Annotated variants lie in 624 unique genes (**Additional File 3**). This leads to an initially high precision of 0.95 (**Fig. 3**), but a modest sensitivity of 0.14.

**Fig. 3.**
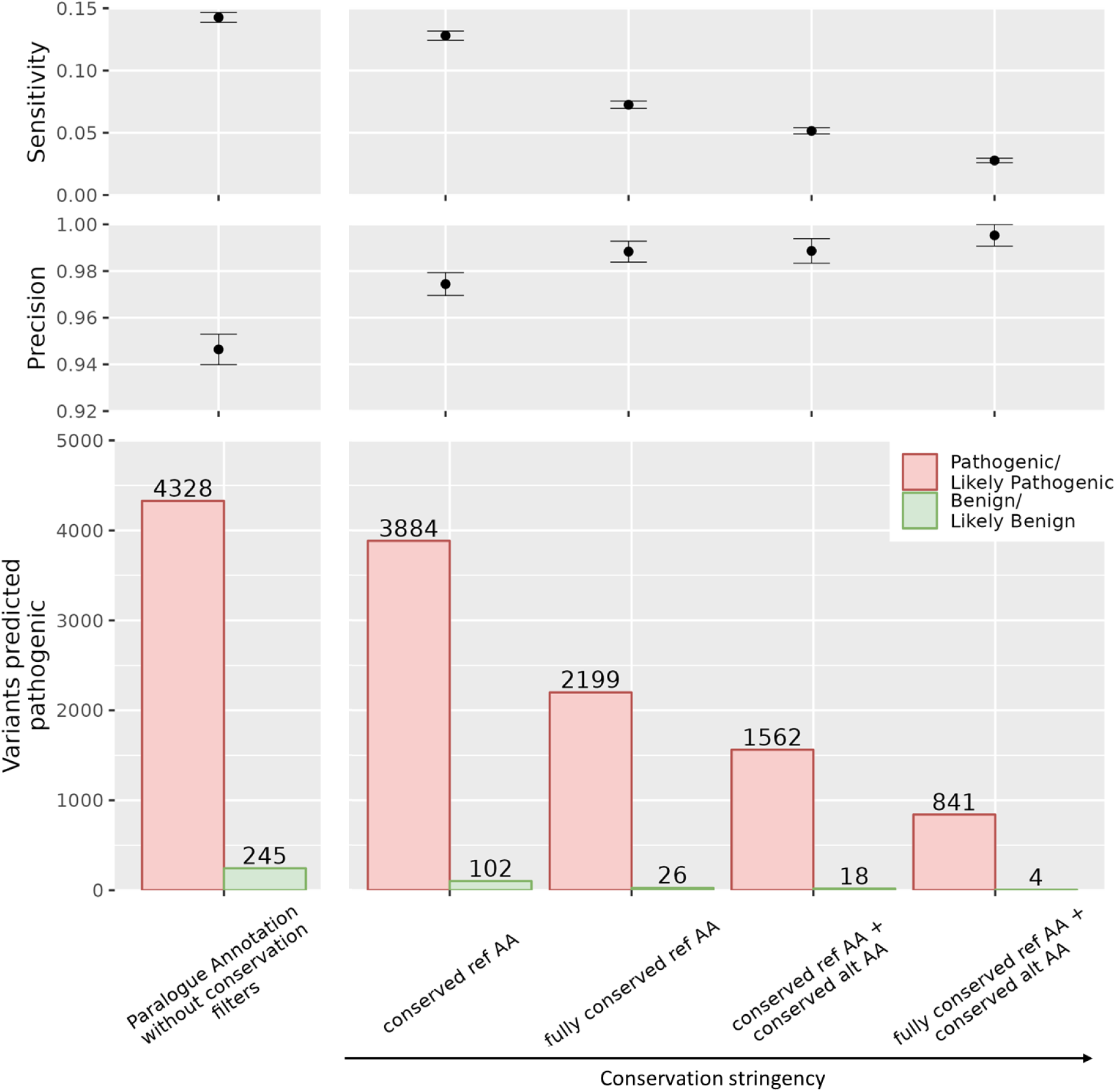
Paralogue annotation has high precision. 95% of variants annotated as pathogenic using this approach are true positives. Variants at positions with increasing conservation stringency (i.e. conservation of the reference and/or alternate amino acid between the query variant and paralogue variant) are even more likely to be pathogenic. The histogram shows the number of P/LP variants (red) and B/LB variants(green) that are annotated as pathogenic by paralogue annotation natively, and with increasingly stringent conservation filters applied, with precision and sensitivity shown above.

Increasing alignment stringency, as a measure of confidence that two aligned amino acids are “equivalent”, leads to improved precision. If we require the query gene and paralogue to have the same reference amino acid at the position of interest, 3,884 P/LP and 102 B/LB variants are predicted to be pathogenic (precision = 0.97), with modest reduction in sensitivity (0.13). Increasing stringency further, by considering only positions where the reference amino acid is fully conserved across the whole protein-family, increases precision to 0.99, but with a substantial decrease in sensitivity to 0.07. If we also require identity of the variant allele when transferring annotations the precision approaches 1.0, but with sensitivity of only 0.03 (**Table S1, Fig. 3**).

Note, however, that there are many true positives amongst the variants that do not reach these more stringent requirements. To further evaluate the optimum balance between precision and sensitivity, we interrogated the variants that would be *removed* by each filter of increasing stringency, and calculated the proportion of each discarded bin that was a true positive. Paralogue annotations with non-conserved reference amino acids had the lowest precision (0.76), and most of the *FP* benign variants were filtered by this step (143 out of the total 241 B/LB removed). At all subsequent stages the discarded variants were predominantly true positive with precision amongst the filtered variants of 0.96 or above. In subsequent analyses we therefore required conservation of the ref allele between the query and source variant in order to transfer annotations, but do not discard annotations that do not meet more stringent filtering thresholds considering that a modest further increase in precision does not justify the loss of sensitivity. However variants that do meet these enhanced criteria can be considered with even greater confidence.

A quantitative measure of conservation across the protein family (the Para Z score^19^) provides an alternative approach to define conservation and hence residue equivalence, with more granularity than the simple categorical grouping explored above. We observe consistent results, with a monotonic trade-off between precision and sensitivity with increasing Para Z score (**Fig. 4**), with values intermediate between our conserved ref AA and fully conserved ref AA categorical bins. This provides us with more refined estimates of the probability that an annotated variant is truly pathogenic across the full spectrum of conservation contexts.

**Fig. 4.**
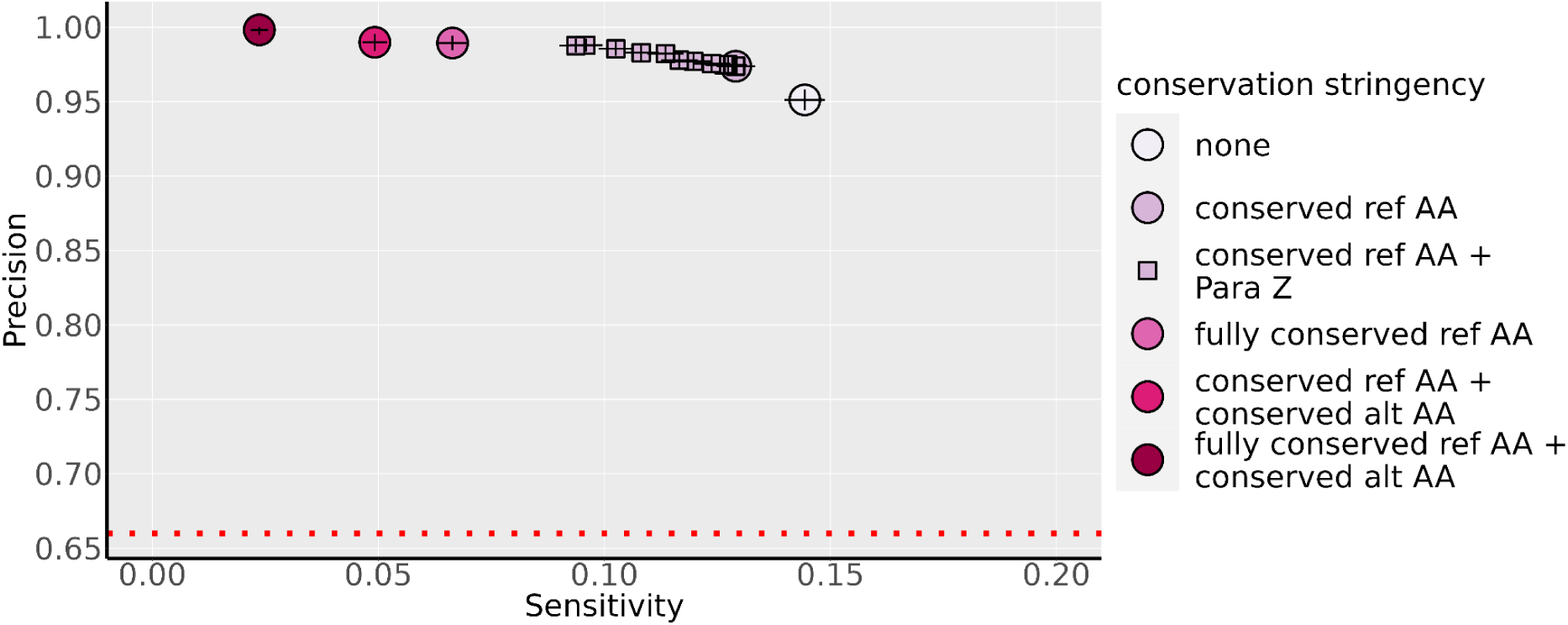
Precision is improved by restricting paralogue annotation to residues that show some degree of conservation across the protein family. Performance of PA is shown alone, and with increasingly stringent conservation filters: requiring identity of the reference amino acid (ref AA) between query & reference variant; requiring identity of the ref AA across the whole protein family (“fully conserved ref AA”); and/or requiring identity of the variant amino acid (alt AA). The ParaZ score is a quantitative measure of conservation across the protein family, ranging from 0-11, with 11 equivalent to a fully conserved ref AA, and lower values representing intermediate degrees of conservation (see Methods Fig. 1). The red dotted line represents baseline precision if predictions were random (determined by the ratio of pathogenic:benign variants in each test set).

To dissect contributions to the high precision achieved by PA in combination with conservation stringency filters, we evaluated the performance of conservation across the protein family as a stand-alone predictor of pathogenicity (**Table S2**). The presence of an aligned residue in a paralogue does not have predictive value in isolation (precision 0.65 demonstrates chance performance in a test set comprising 65% P/LP variants). Variants at positions that are conserved across the protein family are more likely to be pathogenic, with a precision reaching 0.92 at residues that are fully conserved across the whole protein family (8,391 P/LP; 700 B/LB), but the precision is lower than for PA. Finally, to rigorously assess the performance of PA independent of conservation we created a new balanced test set by randomly sampling 700 P/LP variants observed at fully-conserved positions alongside 700 B/LB variants also observed at fully-conserved positions. With one thousand permutations of the P/LP sample the average precision was 0.87, with sensitivity 0.26 (**Fig S2 and S3**). PA has high precision both alone and in combination with conservation stringency filters.

To examine whether the performance of PA depends on the size of the paralogue family, we have conducted an additional sensitivity analysis using the baseline PA (without conservation filters) applied on ClinVar variants. We stratified all protein-coding genes according to their number of paralogues into three groups: small (quantile 0∼33%: 1-3 paralogues), medium (quantile 33%-67%: 4-15 paralogues) and large (quantile 67%-100%:>=16). We found that across three groups, the PPVs are very close to each other (**Table S3**). In contrast, sensitivity was higher for genes with a larger number of paralogues. This is expected, as variants in genes from larger paralogue families are more likely to have paralogous positions and therefore more likely to align with a ClinVar P/LP variant. Overall, paralogue family size influences sensitivity but not PPV.

### Paralogue annotation has a higher positive predictive value than widely-used *in silico* prediction tools

We consider paralogue annotation distinct from typical *in silico* prediction tools, in that it is reporting the presence of a variant that has been previously *directly observed and characterised* as pathogenic for human disease, albeit in an equivalent position of a related protein rather than in the protein of interest. While the presence of such a variant is predictive of pathogenicity, the absence of such a variant does not imply benignity, at least at the present time when most possible human variants have not yet been characterised - this is absence of evidence for pathogenicity, rather than evidence of absence of pathogenicity.

Notwithstanding this, we compared the performance of PA as a binary classifier against widely used *in silico* tools, including AlphaMissense^20,21^, EVE^22^, REVEL ^23^, SIFT^24^, M-CAP ^25^ and CADD ^25^. AlphaMissense and EVE showed higher precision than baseline PA (without conservation filters). With conservation filters applied, PA achieved even higher precision than these two methods. PA showed higher precision than M-CAP, REVEL, CADD, and SIFT across multiple genes and test sets (**Fig. 5A**). PA maintained higher precision than other methods even at relaxed conservation stringency (precision = 0.97), reaching a maximum precision of 0.997 at full conservation. However, PA showed relatively low sensitivity, as many variants have not yet been observed and characterised in humans. Currently, all other methods showed sensitivities of at least 0.89 for identifying P/LP variants in ClinVar (**Fig. 5A**), whereas PA ranged from 0.03–0.16. However, sensitivity is expected to increase as the number of characterised variants available for PA grows. Notably, for specific genes with many disease-associated paralogues, such as SCN5A, sensitivity was higher (0.52) without loss of precision (**Fig. 5B**).

**Fig. 5.**
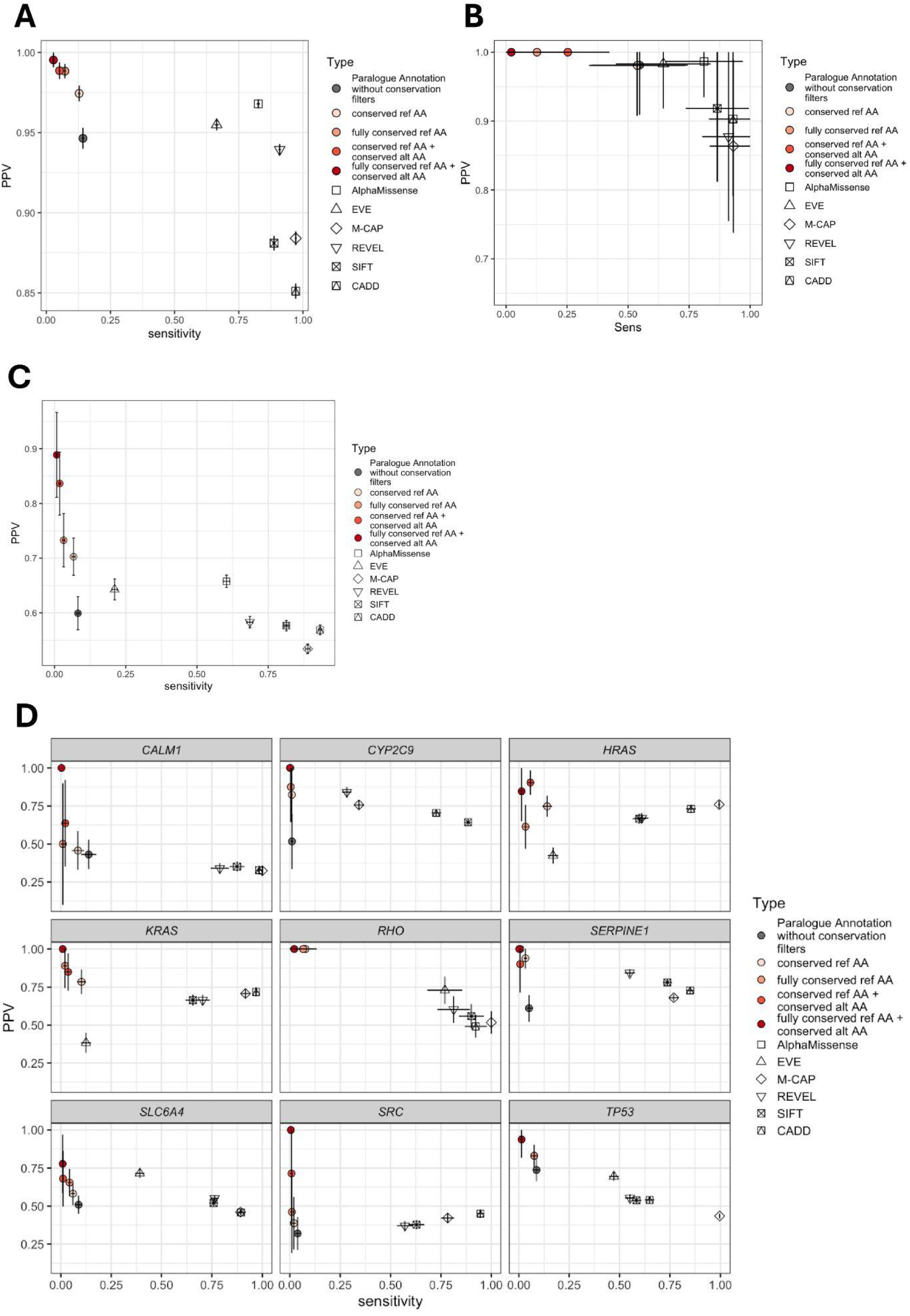
Paralogue Annotation has greater precision than widely used in silico variant prioritisation tools, albeit with limited sensitivity. The precision and sensitivity of Paralogue Annotation (with varying levels of conservation stringency) is compared against the performance of AlphaMissense, EVE, SIFT, M-CAP, CADD and REVEL with each tool applied to the same four sets of variants: **A)** the whole ClinVar dataset; **B)** ClinVar variants in SCN5A; **C)** variants with MAVE fitness measures ; and **D)** variants in the nine genes with MAVE fitness measures assessable with all the benchmarked methods. Error bars represent 95% confidence intervals. In each panel the red dotted line represents baseline precision if predictions were random (determined by the ratio of pathogenic:benign variants in each test set). Baselines differ due to a different total number of variants capable of being annotated by each tool.

While ClinVar is probably the most comprehensive reference dataset of variants with a characterised role in human disease, and spans a wide range of different diseases, it has many inherent biases. We therefore performed a series of additional validations using functional data with multiplexed assays of variant effects (MAVE) available through ProteinGym^26^. In total, 64 genes with both MAVE assays and the baseline paralogue annotation (without conservation filters) were included. Across the MAVE datasets, we found the overall precision of baseline paralogue annotation was comparable to that of some established methods, including SIFT, REVEL, CADD and M-CAP. Increasing conservation stringency by requiring conserved reference AAs could further increase the precision, exceeding that of AlphaMissense and EVE (**Fig. 5C**).

Among the 64 genes, there are nine that we can evaluate at the individual gene level using all the methods (**Fig.5D**). For genes *CALM1*, *HRAS*, *KRAS*, *RHO*, and *TP53*, the baseline paralogue annotation has comparable performance or even better than the established methods, highlighting the value of applying PA to these gene families. For other genes, although the baseline PA is inferior to the established VEP methods, increasing conservation stringency can improve the precision, even to a level higher than that of those methods. Note that the balance of each test set was different, so the precisions are not comparable between test sets, but are comparable between tools within each test set.

### Additional validation approaches not reliant on gold-standard classifications confirm that paralogue annotation identifies disease-associated variants

We performed additional performance assessments independent of classification labels using human case-control data from two cohorts with inherited arrhythmia syndromes (Long QT syndrome (LQTS) and Brugada syndrome (BrS)) and a cohort with neurodevelopmental disorders with epilepsy (NDD+E). We evaluated associations between rare variants in established disease genes and inherited diseases without relying on pathogenicity assertions for individual variants. Disease associations were assessed across five variant groups: (i) rare missense variants alone; (ii) rare variants occurring at conserved paralogous sites; (iii) rare variants prioritised by paralogue annotation; and (iv–v) variants meeting both PA and paralogue conservation criteria, defined as variants with either a conserved RefAA (iv) or a fully family-wise conserved RefAA (v).

Rare variants prioritised by paralogue annotation (PA) alone showed stronger disease associations than rare variants alone. For two genes involved in LQTS – *KCNQ1* and *KCNH2* (**Fig. 6**), PA priortised variants showed substantially stronger disease associations than unselected rare missense variants (OR 162 vs 60; and 103 vs 34). For *SCN5A*, PA prioritised variants do not show significantly stronger association with LQTS (OR 8 vs 5), but do show a stronger association with BrS (OR 54 vs 30), possibly reflecting different molecular mechanisms underlying these diseases (Loss-of-function in BrS vs gain-of-function in LQTS). For genes implicated in NDD+E (SCN1A, SCN8A, and KCNQ2), applying PA resulted in biologically and statistically significant increases in odds ratios, from a mean of 1.2 across the three genes to 12.7. In contrast, no significant difference was observed for SCN2A, where infantile-onset epilepsy is understood to be caused by gain-of-function variants, while loss-of-function variants are associated with autism spectrum disorder (ASD) or intellectual disability without epilepsy, or with mild or later-onset epilepsy^37,38^.

**Fig. 6.**
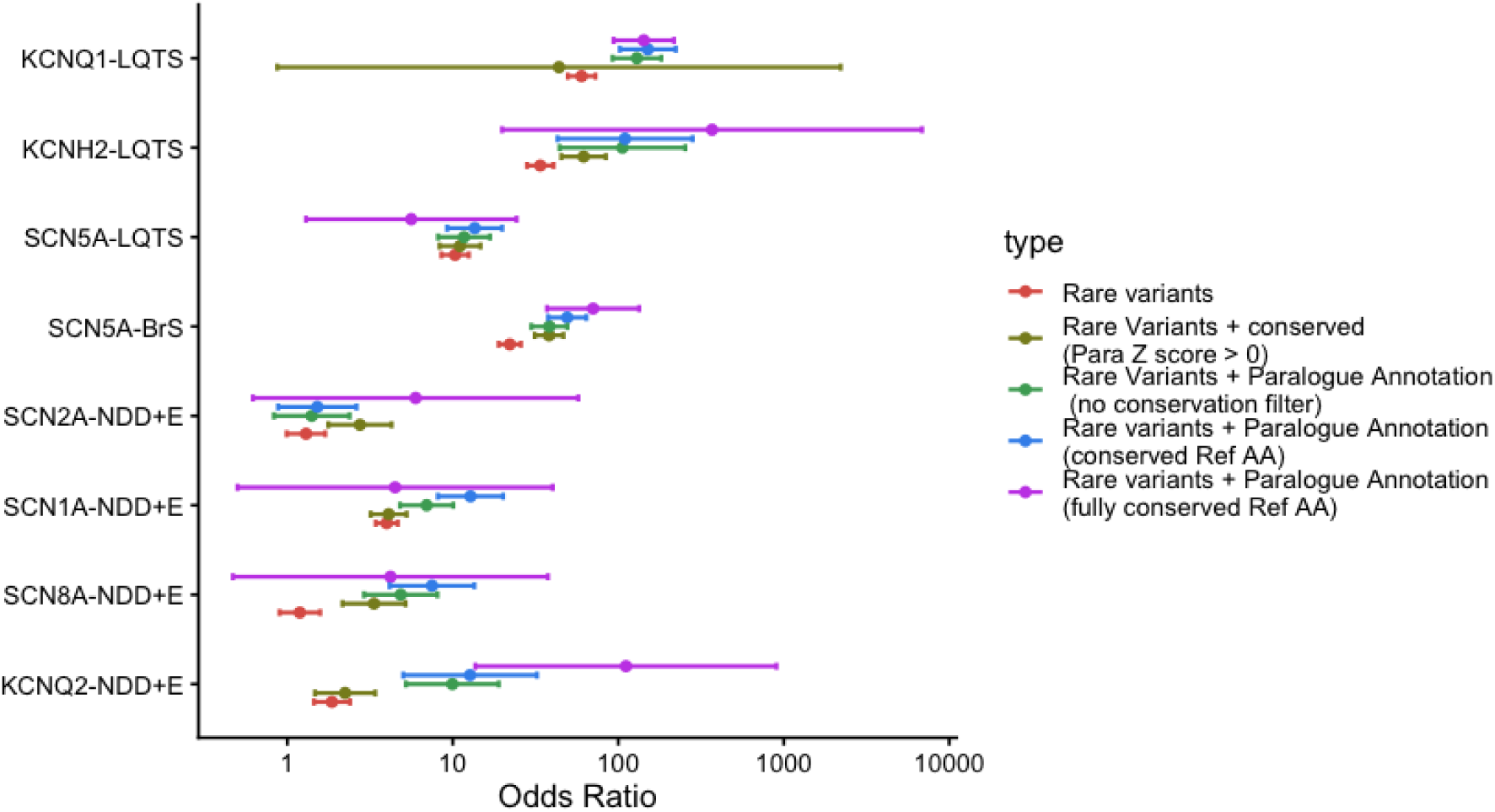
Variants prioritised by paralogue annotation show stronger disease associated than unstratified rare variants. Shown are Odds Ratios calculated from case and control series for: KCNQ1, KCNH2 and SCN5A in Long QT Syndrome (LQTS); SCN5A in Brugada Syndrome (BrS); SCN2A, SCN8A, SCN1A, and KCNQ2 in Neurodevelopmental Disorder with Epilepsy (NDD+E). Odds ratios are presented for the following variant groups: all rare variants (red); rare variants at paralogue-conserved sites (olive, Para Z score > 0); rare variants predicted to be pathogenic by paralogue annotation (green, without conservation filters); rare variants with paralogue annotation at conserved reference AA (blue); and rare variants with paralogue annotation at fully conserved reference AAs (purple). Error bars show 95% confidence intervals. For SCN5A-LQTS and SCN2A-NDD+E, the mechanism of disease is understood to be Gain-of-Function, while other genes cause disease by Loss-of-Function.

Consistent with previous observations that disease-associated missense variants are enriched at paralogue-conserved sites^19^, rare variants occurring at conserved paralogue sites also showed stronger disease associations than rare variants alone. Across several gene–disease pairs, PA achieved higher odds ratios than paralogous conservation alone, including *KCNQ1*, *KCNH2*, and *SCN5A* in long QT syndrome, and *SCN1A*, *SCN8A*, and *KCNQ2* in neurodevelopmental disorder with epilepsy. Combining PA with paralogous conservation (using a conserved RefAA or a fully conserved RefAA) further increased odds ratios compared with either criterion alone.

Together, these results demonstrate that PA alone meaningfully improves variant prioritisation, and that its performance can be further enhanced when combined with site conservation.

Finally, we studied *de novo mutations (DNM)* in individuals with and without ASD. Individuals with ASD have been previously shown to have a higher burden of “deleterious” missense variants exome-wide, where deleteriousness is defined by *in silico* predictors. We hypothesised that paralogue annotation would likewise stratify a pool of variants that are enriched in disease. DNM annotated as pathogenic by PA were more prevalent in those with ASD than controls. While PA prioritised far fewer variants than *in silico* predictors (38 and 14 DNM from cases and controls respectively for PA compared to 431 and 233 for REVEL, which prioritised the least out of all the predictors), PA-annotated variants were much more strongly enriched (**Table 1**). This enrichment is statistically significant compared to all missense DNM across a range of PA conservation stringencies (none; conserved ref AA; or fully conserved ref AA) (**Table S4**).

**Table 1.**
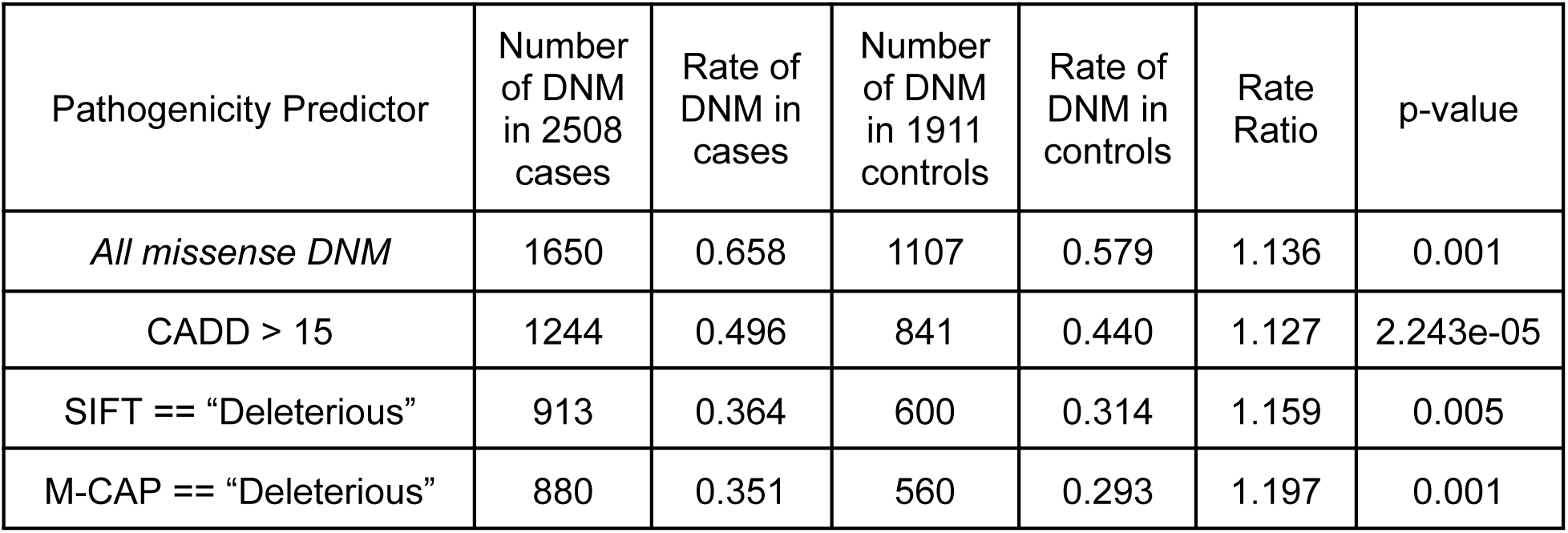
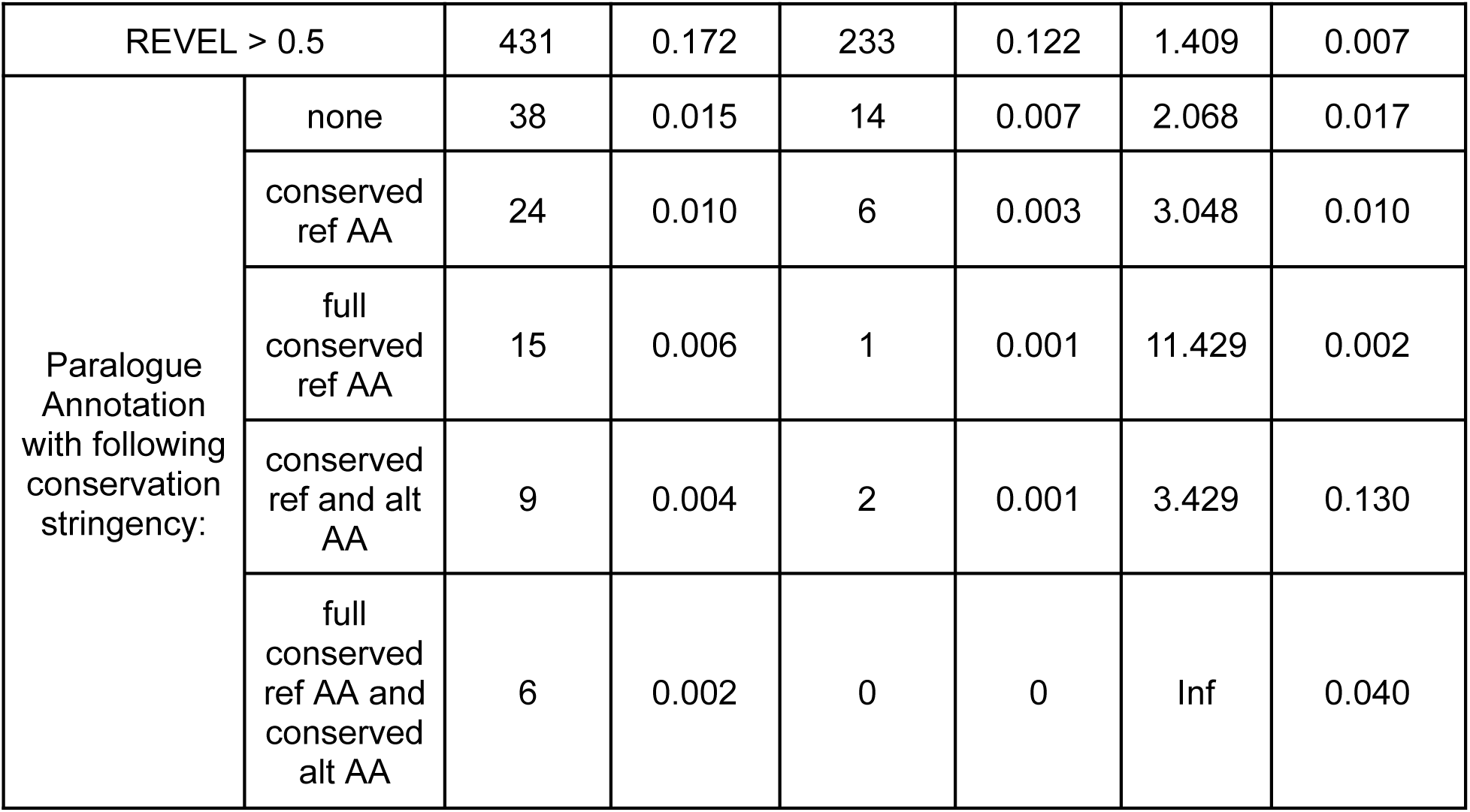
De Novo Mutations (DNM) annotated as pathogenic by Paralogue Annotation are highly enriched in individuals with Autism Spectrum Disorder (ASD) patients. Shown are all DNM missense variants found by trio exome sequencing from 2508 ASD patients and 1911 unaffected siblings, stratified by in silico pathogenicity predictors and Paralogue Annotation at varying conservation stringencies. ASD patients have a higher rate of de novo missense variants than controls.

| Pathogenicity Predictor | Number of DNM in 2508 cases | Rate of DNM in cases | Number of DNM in 1911 controls | Rate of DNM in controls | Rate Ratio | p-value |
| --- | --- | --- | --- | --- | --- | --- |
| *All missense DNM* | 1650 | 0.658 | 1107 | 0.579 | 1.136 | 0.001 |
| CADD > 15 | 1244 | 0.496 | 841 | 0.440 | 1.127 | 2.243e-05 |
| SIFT == “Deleterious” | 913 | 0.364 | 600 | 0.314 | 1.159 | 0.005 |
| M-CAP == “Deleterious” | 880 | 0.351 | 560 | 0.293 | 1.197 | 0.001 |

|  |  |  |  |  |  |  |  |
| --- | --- | --- | --- | --- | --- | --- | --- |
| REVEL > 0.5 |  | 431 | 0.172 | 233 | 0.122 | 1.409 | 0.007 |
| Paralogue Annotation with following conservation stringency: | none | 38 | 0.015 | 14 | 0.007 | 2.068 | 0.017 |
|  | conserved ref AA | 24 | 0.010 | 6 | 0.003 | 3.048 | 0.010 |
|  | full conserved ref AA | 15 | 0.006 | 1 | 0.001 | 11.429 | 0.002 |
|  | conserved ref and alt AA | 9 | 0.004 | 2 | 0.001 | 3.429 | 0.130 |
|  | full conserved ref AA and conserved alt AA | 6 | 0.002 | 0 | 0 | Inf | 0.040 |

### Extending this approach to transfer annotations across homologous protein domains (Pfam) retains high precision and increases the number of informative annotations

While 14,514 protein-coding genes have a closely related paralogue, an even larger number contain protein domains that have homologues elsewhere in the genome. We sought to evaluate whether the same principles could be applied across homologous protein domains shared between genes that are *not* paralogues. 18,083 genes contain at least one Pfam domain, with 13,537 genes both having at least one paralogue and also containing at least one Pfam domain. Importantly, 4,546 genes without paralogues do nonetheless contain Pfam domains, which might yield informative annotations (**Fig 7a**).

**Fig. 7.**
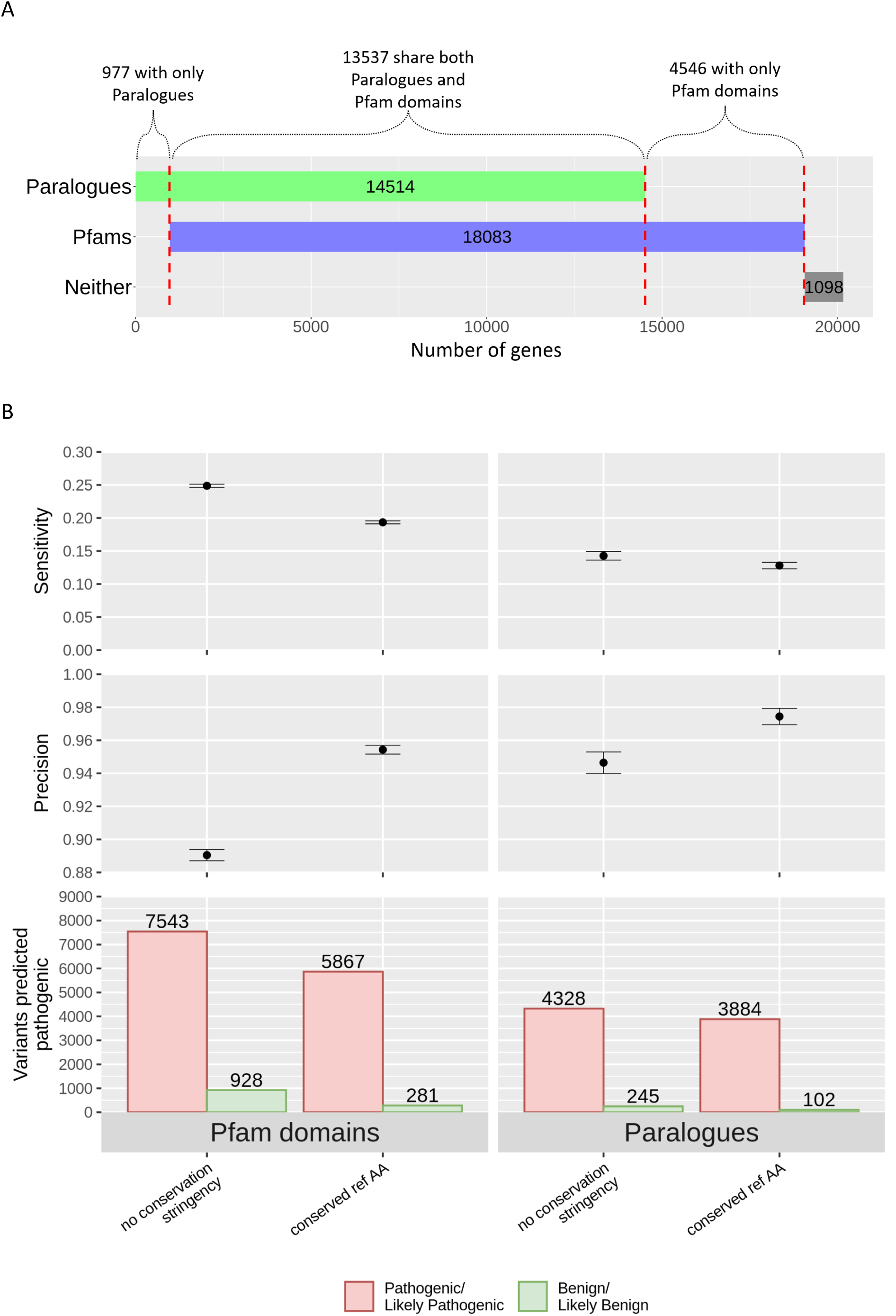
Transferring annotations across homologous residues in Pfam domains increases the number of annotations, and maintains high precision. **(A) There are more genes that contain at least one Pfam domain than have a paralogue.** The number of genes that have at least one paralogue is shown in green, and the number with at least one Pfam domain in blue. Genes with neither are shown in grey. **(B) Transferring annotations between homologous Pfam domains yields more annotations than Paralogue Annotation, while maintaining high precision.** There are more Pathogenic/Likely Pathogenic (red) and Benign/Likely Benign (green) predictions even when both query and aligned variants are conserved.

Pfam “metadomain” alignments^36^ identify homologous positions in Pfam domains across different proteins, and provide a convenient method to extend this approach. Extending our approach from paralogues to Pfam metadomains more than doubles the number of variants annotated (from 4,328 to 7,543 for P/LP variants). There is a also a larger proportion of known B/LB variants receiving a false positive paralogue annotation (245 to 928) (**Fig 7b**), yielding a 6% absolute reduction in precision (0.95 to 0.89) (**Table S5**), but with an important 74% increase in sensitivity (0.14 to 0.25). As before, precision is improved by considering only variants at positions with conserved reference amino acids, maintaining precision at 0.95 while still increasing sensitivity (0.19) over PA (**Fig 7b; Table S6**).

### Paralogue Annotation annotates many variants of uncertain significance, and could potentially annotate ∼10% of all possible missense variants with high precision

In ClinVar, there are 127,080 missense variants classified as VUS, of which 86,894 have an aligned position in a paralogue. Currently, 3,508 can be annotated by at least one known P/LP at an equivalent position of a paralogue (**Table 4**). This provides evidence in support of pathogenicity for 5% of missense VUS from disease genes with disease paralogues (3% of total VUS).

**Table 4.**
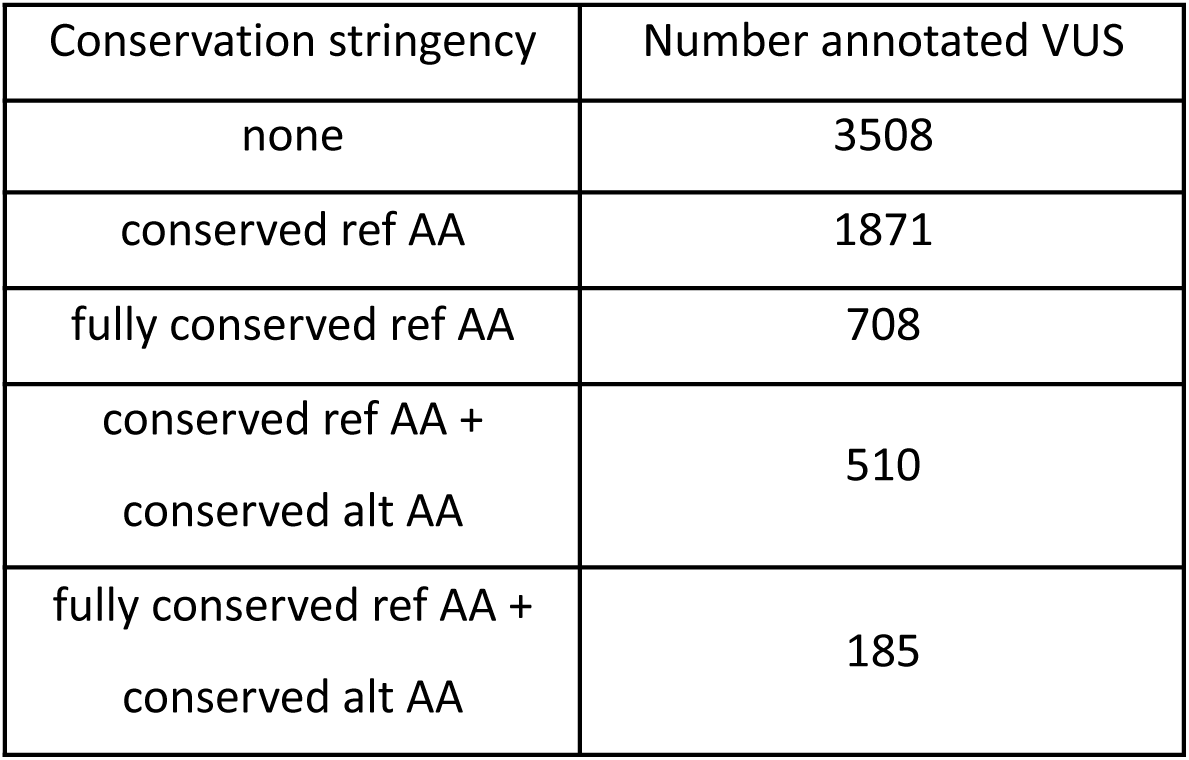
Number of ClinVar VUS predicted pathogenic by Paralogue Annotation. Shown are missense variants of uncertain significance (VUS) from ClinVar that are predicted to be pathogenic by Paralogue Annotation at varying conservation stringencies.

Considering all possible single nucleotide changes that could result in a missense variant within the human GRCh37 reference genome (179,684,761), we identified that 20,971,151 of these variants occur at a position with an equivalent residue in an aligned paralogue. With the 30,334 ClinVar variants used in this study, 855,552 additional missense variants (at 335,243 distinct genomic positions) could be annotated by a known Pathogenic/Likely pathogenic in a paralogue.

Considering positions at which reference alleles are conserved, and where PA yields higher confidence annotations, 466,892 additional possible missense variants (at 176,889 genomic positions) could be annotated. These numbers will increase as the number of well-characterised variants deposited in ClinVar, or equivalent reference resources, increases.

### Variants predicted as pathogenic by Paralogue Annotation or annotation across homologous Pfam domains are available through a user-friendly webtool

In addition to command line tools to perform paralogue annotation, and code to replicate these analyses, we have developed an online webtool (available at https://www.cardiodb.org/paralog_app) that does not require users to download or install the software used in this study, to ensure that Paralogue Annotation is readily accessible to the wider community.

## Discussion

While traditional homology based bioinformatic methods, like SIFT, have utilized alignments of conserved sequences to assess pathogenicity, PA takes this a step further by directly considering information about human variation. This is done by transferring annotations of variant consequences between evolutionarily-related proteins and domains, with variant effect evaluated by directly observing the consequence of variation in humans on disease phenotypes.

The approach is widely applicable. The majority of human disease-associated genes across diverse protein classes have a paralogue and/or a shared Pfam domain, and a large proportion of these (54%) contain variants annotated with human disease consequence. The potential for informing variant interpretation is therefore substantial: the 30,334 P/LP missense variants analysed here can annotate and inform interpretation of a further 855,552 possible missense variants across the genome, representing a 30-fold increase. And while we use ClinVar as the most comprehensive curated public collection of human variants annotated with known disease consequence, the method can be applied to any set of variants.

Paralogue Annotation stands out for its high positive predictive value. This is dependent on the stringency of the assertion that two amino acid substitutions in related proteins or related domains are “equivalent”. This is highest when both the reference amino acid and substituting amino acid are identical, but precision remains high even when different substitutions are seen at a conserved position. For most purposes we recommend applying paralogue annotation if there is pairwise conservation of the reference amino acid between the two genes under consideration. Although more stringent filters improved precision, the majority of variants eliminated were true positives. We anticipate that there is further opportunity for refinement here - for example considering physicochemical similarity of reference and alternate alleles rather than solely AA identity - which may improve precision without such a marked sensitivity penalty. It is also important to consider equivalence of disease-consequence. There is insufficient structured data to allow a systematic analysis of PA stratified by disease mechanism (gain vs loss of function), but we would anticipate annotations to be most information where a pair of disease-associated genes have concordant pathogenic mechanisms. We recommend users with a specific disease gene of interest to first check if pathogenic variants found in paralogues function by the same disease mechanism prior to making a final judgement on annotations identified.

PA can be readily incorporated into the existing ACMG/AMP framework. Criteria PS1 (same amino acid change as an established pathogenic variant) and PM5 (novel missense variant at a residue position where a different missense substitution is established as pathogenic) can be applied to missense variants observed at equivalent positions across paralogues. The ClinGen RASopathy Expert Panel have adopted this approach^10^, and apply the rule at the same level of strength for a variant seen in the query gene or specific very closely related paralogues. Other working groups are piloting the approach by applying the rule at a lower level of evidence for variants in paralogues (e.g. PS1-moderate, PM5-supporting)(personal communication).

The principal apparent limitation of PA alongside other tools is its relatively low sensitivity. There are several explanations for this. Fundamentally, this approach can never be applied to a gene that has no paralogues, and cannot be applied to a region that is divergent between paralogues without 1:1 equivalence in a sequence alignment. The first limitation is partially addressed by transferring annotations between Pfam domains shared beyond direct paralogues, which significantly improves sensitivity while preserving clinically-useful precision. The latter might be addressable by considering other models of equivalence beyond the linear sequence alignment, for e.g. positions that are nearby in 3D protein structures. But there will remain protein regions that do not have orthologous sequence, for example in intrinsically disordered regions.

Furthermore, at present many P/LP variants are not annotated because paralogous genes are not known to be associated with human disease, or simply because variation at the equivalent position in a paralogue has not yet been observed, characterised, and deposited in ClinVar. In these cases PA will become more informative with time, as more disease genes are recognised and annotated as such, and more pathogenic variants in those genes are observed and characterised. For key disease genes, high-throughput functional studies are actively generating comprehensive datasets of functionally-charracterised variants^39^, which may also be used to annotate across paralogues. Nonetheless, a limited sensitivity does not imply a lack of clinical value. ClinVar annotates approximately 1.2% (855K out of 70M all possible missense variants) of possible missense variants in the genome as either P/LP or B/LB. The sensitivity of ClinVar for the identification of pathogenic variants is unknown, but is presumably very low. Yet ClinVar remains a first line resource because the information is extremely valuable for the small number of variants annotated. The high precision of PA indicates that it provides useful evidence when a variant is annotated.

Finally, we note the absence of a paralogue annotation should be treated as an absence of evidence for pathogenicity, but this does not provide evidence of benignity. As data accumulates, the approach might be extended to consider transferring assertions of benignity, but this comes with additional reservations as laboratories may have widely differing practice for reporting and depositing variants that are benign with respect to the phenotype observed in a patient undergoing testing, but may not be benign with respect to a different phenotype.

## Conclusions

Transferring pathogenicity annotations between equivalent substitutions in human paralogues, or across homologous Pfam protein domains, identifies pathogenic variants with high precision. This approach is broadly applicable across a diverse range of human disease genes. In the context of clinical variant interpretation precision is arguably the most important performance metric; since an incorrect genetic diagnosis has potential to cause greater harm than a false negative genetic test, clinical genetic diagnostics usually prioritises control of the false positive rate over the false negative rate. Although sensitivity is currently limited, this is in large part due to sparse datasets of well-characterised variants in resources such as ClinVar, and such reference resources are growing rapidly with increasing availability of clinical sequencing, such that the utility of these approaches will increase.

If abbreviations are used in the text they should be defined in the text at first use, and a list of abbreviations should be provided.

## List of abbreviations

AA: Amino Acids
ACMG/AMP: American College of Medical Genetics and Genomics/Association for Molecular Pathology
alt: Alternate (allele)
ASD: Autism Spectrum Disorder
B/LB: Benign/Likely benign
BrS: Brugada Syndrome
DNM: *De Novo* Mutations
LQTS: Long QT Syndrome
MSA: Multiple Sequence Alignment
OR: Odds Ratio
PA: Paralogue Annotation
P/LP: Pathogenic/Likely pathogenic
ref: Reference (allele)

## Declarations

### Ethics approval and consent to participate

The data used in this analysis has been previously published and/or is in the public domain, and the study did not use identifiable or sensitive data. No additional human samples or data were collected for this study. Therefore UK research ethics committee approval was not required.

### Consent for publication

Not applicable

### Availability of data and materials

The locations of publicly available datasets analysed during the current study are cited in the methods.

All new data generated during this study are included in this published article [and its supplementary information files].

Code to reproduce the analyses are available at https://github.com/ImperialCardioGenetics/ParalogueAnnotation_supplementary_repo.

The paralogue annotation VEP plug-in is available at https://github.com/ImperialCardioGenetics/paralogueAnnotator.

The list of 855K all possible missense variants predicted to be pathogenic by Paralogue Annotation, using ClinVar P/LP variants, can be downloaded from our web application at https://www.cardiodb.org/paralog_app/.

To query the paralogous annotation of a missense variant, or to batch download paralogous annotations for all possible missense variants, please use https://www.cardiodb.org/paralog_app/. The code to reproduce this web application is available at https://github.com/ImperialCardioGenetics/Paralogue_Annotation_App.

## Authors’ contributions

NL, RW, NW & JSW conceived and designed the analysis; NL, XZ & RW collected the data; EM, PT, MJ, XZ, MA, GP, RW, HH & DL contributed data or analysis tools; NL & XZ performed the analysis; NL, XZ & JSW wrote the paper. All authors reviewed the manuscript for important intellectual content.

## Acknowledgements

We thank Laurens Wiel and the developers of metadome for providing the Pfam domain alignments used in this study. We also thank Emily Perry, Fiona Cunningham and the Ensembl Genome Interpretation Team for the support they provided during the development of the Paralogue Annotation plugin for VEP.

