## Supplementary Material for "Variant annotation across homologous proteins (“Paralogue Annotation”) identifies disease-causing missense variants with high precision, and is widely applicable across protein families"

**Supplementary tables, figures and additional files for "Variant annotation across homologous proteins ("Paralogue Annotation") identifies disease-causing missense variants with high precision on an exome-wide scale"**

***Table of contents***

***Table S1) Number of ClinVar variants predicted to be pathogenic using Paralogue Annotation.***
***Table S2) Number of aligned paralogous positions at varying paralogous conservation among ClinVar variants.***
***Table S3) Performance of Paralogue annotation stratified by the size of paralogue family (number of paralogues).***
***Table S4) 95% Confidence Intervals from bootstrap distribution of differences in rate ratios from Table 1 .***
***Table S5) Precision (Positive Predictive Value - PPV) decreases marginally when using Pfam metadomain alignments instead of paralogue alignments within the Paralogue Annotation framework***
***Table S6) Sensitivity increases substantially when using Pfam metadomain alignments instead of paralogue alignments within the Paralogue Annotation framework.***

***Figure S1) Precision of paralogue annotation on a balanced test set of 700 P/LP and B/LB ClinVar variants at positions where the reference reference amino acid is fully conserved across the paralogue family (1000 samples).***
***Figure S2) Sensitivity of balanced 700 P/LP and B/LB ClinVar variants with fully conserved reference amino acids across 1000 repeats .***

| | | Pathogenic variants | | Benign variants | | PPV | Sensitivity | P-value |
|---|---|---|---|---|---|---|---|---|
| Total variants | | 30334 | | 16682 | | *0.65** | - | - |
| Test | | Test Positive (TP) | Test Negative (FN) | Test Positive (FP) | Test Negative (TN) | | | |
| (a) Does variant align to a P/LP paralogue variant? | | 4328 | 26006 | 245 | 16437 | 0.95 | 0.14 | <5.0e-324 |
| increasing conservation stringency | (b) Does variant align to a P/LP paralogue variant with the same conserved ref AA? | 3884 | 26450 | 102 | 16580 | 0.97 | 0.13 | <5.0e-324 |
| | (c) Does variant align to a P/LP paralogue variant where ref AA is fully conserved family-wide? | 2199 | 28135 | 26 | 16656 | 0.99 | 0.07 | <5.0e-324 |
| | (d) Does variant align to a P/LP paralogue variant with the same conserved ref and alt AA? | 1562 | 28772 | 18 | 16664 | 0.99 | 0.05 | 3.17e-270 |
| | (e) Does variant align to a P/LP paralogue variant where ref AA is fully conserved family-wide and with the same conserved alt AA? | 841 | 29493 | 4 | 16678 | 1.00 | 0.03 | 7.6e-154 |

***Table S1) Number of ClinVar variants predicted to be pathogenic using Paralogue Annotation.*** *Shown are number of ClinVar variants with a clinical significance of Pathogenic/Likely pathogenic (P/LP Variants) and Benign/Likely benign (B/LB Variants) that are predicted to be pathogenic using* ***(a)*** *Paralogue Annotation and the remaining number of variants after filtering at each subsequent conservation stringency level, including:* ***(b)*** *conserved reference amino acids (ref AA);* ***(c)*** *full conserved ref AA;* ***(d)*** *conserved ref AA and conserved alternate amino acids (alt AA); and* ***(e)*** *full conserved ref AA and conserved alt AA. The Precision/Positive Predictive Value (PPV) (* represents baseline PPV if predictions were random), Sensitivity, Specificity and Fisher's Exact test p-value are shown to the nearest 2 significant figures where applicable. Note: machine performing calculations cannot compute values smaller than 5e-324.*

| | Pathogenic variants | | Benign variants | | PPV | Sensitivity | P-value |
|---|---|---|---|---|---|---|---|
| Total variants | 30334 | | 16682 | | *0.65** | | |
| Test | Test Positive (TP) | Test Negative (FN) | Test Positive (FP) | Test Negative (TN) | | | |
| Does variant have an aligned paralogue position? | 23749 | 6585 | 12812 | 3870 | 0.65 | 0.78 | 2.0e-4 |
| Does variant have an aligned paralogue position and ParaZ >= 1? | 15778 | 14556 | 5091 | 11591 | 0.76 | 0.52 | <5.0e-324 |
| Does variant have an aligned paralogue position and ParaZ >= 3? | 15430 | 14904 | 4671 | 12011 | 0.77 | 0.51 | <5.0e-324 |
| Does variant have an aligned paralogue position and ParaZ >= 5? | 14225 | 16109 | 3663 | 13019 | 0.80 | 0.47 | <5.0e-324 |
| Does variant have an aligned paralogue position and ParaZ >= 7? | 12812 | 17522 | 2619 | 14063 | 0.83 | 0.42 | <5.0e-324 |
| Does variant have an aligned paralogue position and ParaZ >= 9? | 10851 | 19483 | 1505 | 15177 | 0.88 | 0.36 | <5.0e-324 |
| Does variant have an aligned paralogue position and ParaZ >= 11? | 9293 | 21041 | 889 | 15793 | 0.91 | 0.31 | <5.0e-324 |
| Does variant have an aligned paralogue position that is fully conserved across the family? | 8391 | 21943 | 700 | 15982 | 0.92 | 0.28 | <5.0e-324 |

***Table S2) Number of aligned paralogous positions at varying paralogue conservations among ClinVar variants.*** *Shown are the number of aligned paralogue positions among ClinVar Pathogenic/Likely pathogenic and Benign/Likely benign variants at increasing ParaZ score thresholds. The number of aligned positions that are also fully conserved across the family are included for comparison.*

| Gene group based on the number of paralogues | Number of genes | PPV | Sensitivity |
|---|---|---|---|
| small | 105 | 94.3% | 5.7% |
| medium | 219 | 92.6% | 11.3% |
| large | 359 | 94.3% | 30.9% |

***Table S3) Performance of Paralogue annotation stratified by the size of paralogue family (number of paralogues).*** *Based on genes' number of paralogues, we classify them into three groups: small (quantile 0~33%: 1-3 paralogues), medium (quantile 33%-67%: 4-15 paralogues) and large (quantile 67%-100%:>16). The Precision (PPV) and Sensitivity of PA are calculated for each group.*

| Paralogue Annotation with following conservation stringency | 95% Confidence Intervals | |
|---|---|---|
| | Lower bound | Upper bound |
| none* | -4.15 | -0.0239 |
| conserved Ref AA* | -13.0 | -0.290 |
| full conserved Ref AA* | -Inf | -2.29 |
| conserved Ref and Alt AA | -Inf | 0.238 |
| full conserved Ref AA and conserved Alt AA | -Inf | -Inf |

***Table S4) 95% Confidence Intervals from bootstrap distribution of differences in rate ratios from Table 1 .*** *Shown are the lower and upper bound 95% Confidence Intervals (CI) calculated from a bootstrap distribution of differences in rate ratios from Table 1 between Paralogue Annotation and All missense DNM. Bootstrap resampling was done under a Poisson distribution using the rates of DNM in cases and controls as the distribution parameter lambda. * indicates CI that do not include 0 and are therefore significant.*

| | PPV | PPV % change | PPV with conserved ref AA | PPV with conserved ref AA % change |
|---|---|---|---|---|
| Pfam | 0.89 | -5.9% | 0.95 | -2.0% |
| Paralogues | 0.95 | | 0.97 | |

***Table S5) Precision (Positive Predictive Value - PPV) decreases marginally when using Pfam metadomain alignments instead of paralogue alignments within the Paralogue Annotation framework.*** *Shown are the precision of Pfam domains compared to paralogues and their percentage change with and without filtering for conserved reference amino acids (ref AA).*

<table>
<tr><th></th><th>Sensitivity</th><th>Sensitivity % change</th><th>Sensitivity with conserved ref AA</th><th>Sensitivity with conserved ref AA % change</th></tr>
<tr><td>Pfam</td><td>0.25</td><td rowspan="2">+74%</td><td>0.19</td><td rowspan="2">+51%</td></tr>
<tr><td>Paralogues</td><td>0.14</td><td>0.13</td></tr>
</table>

***Table S6) Sensitivity increases substantially when using Pfam metadomain alignments instead of paralogue alignments within the Paralogue Annotation framework.*** *Shown are the sensitivity of Pfam domains compared to paralogues and their percentage change with and without filtering for conserved reference amino acids (ref AA).*

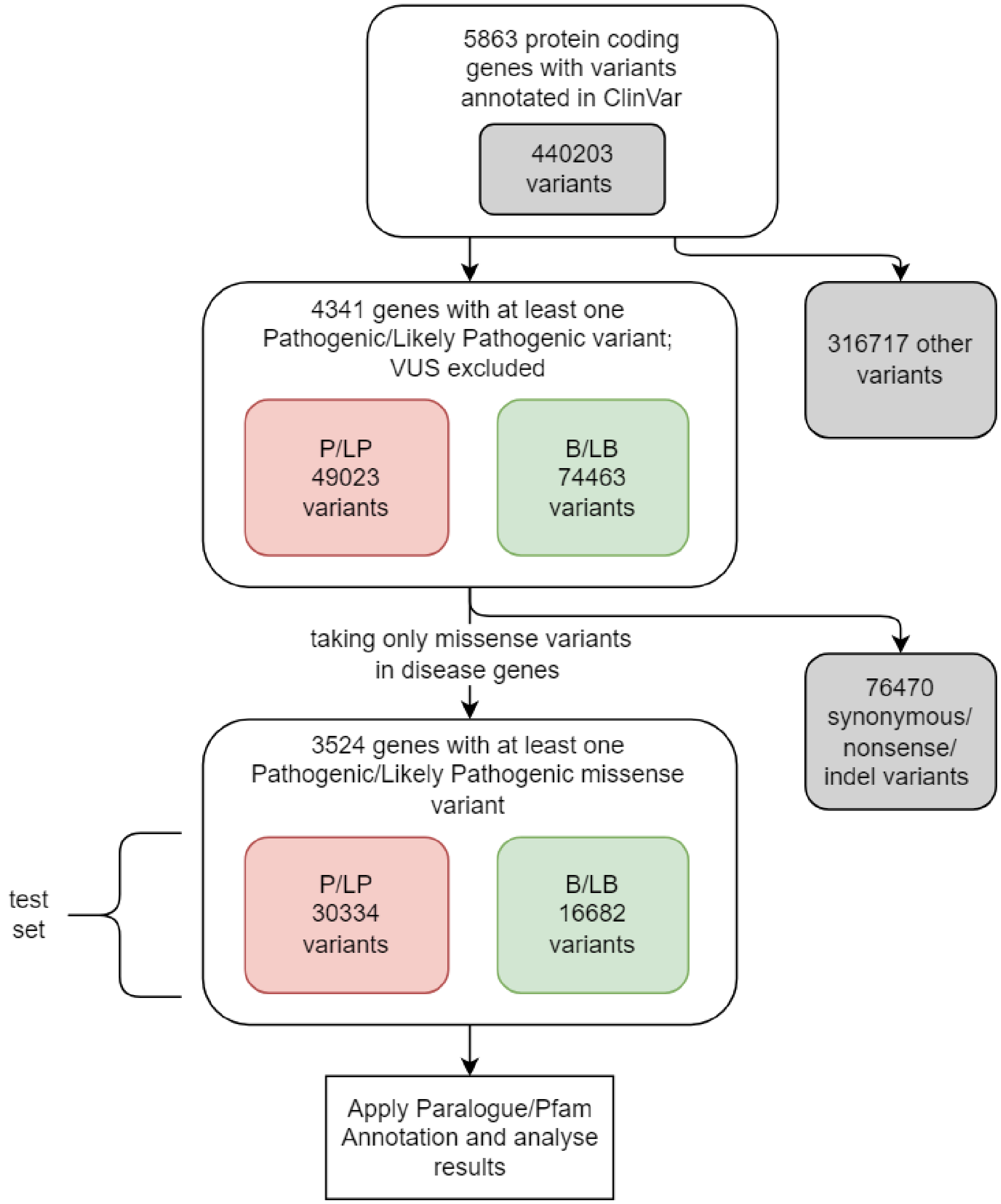


***Figure S1) Outline of experimental design and selection of ClinVar variants for annotation.***

Variants were drawn from ClinVar version 20190114 (build GRCh37) .

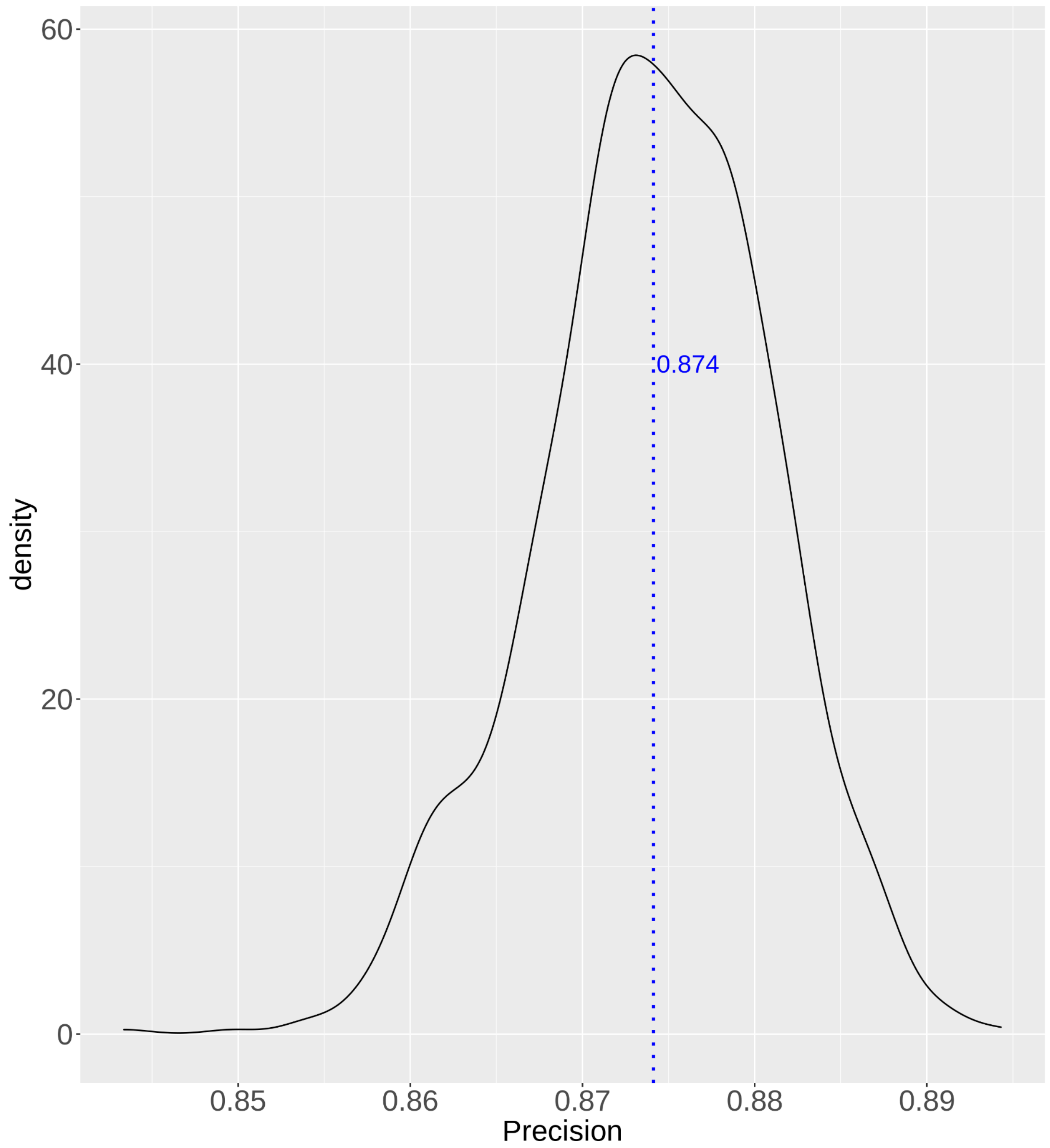


***Figure S2) Precision of paralogue annotation on a balanced test set of 700 P/LP and B/LB ClinVar variants at positions where the reference amino acid is fully conserved across the paralogue family (1000 samples).*** *Shown is a kernel density estimation plot of precisions calculated from 1000 repeated tests of a balanced set of 700 randomly sampled Pathogenic/Likely pathogenic (P/LP) variants and 700 Benign/Likely benign (B/LB) variants at positions that are fully conserved across the family. Average precision is shown by the blue dotted line.*

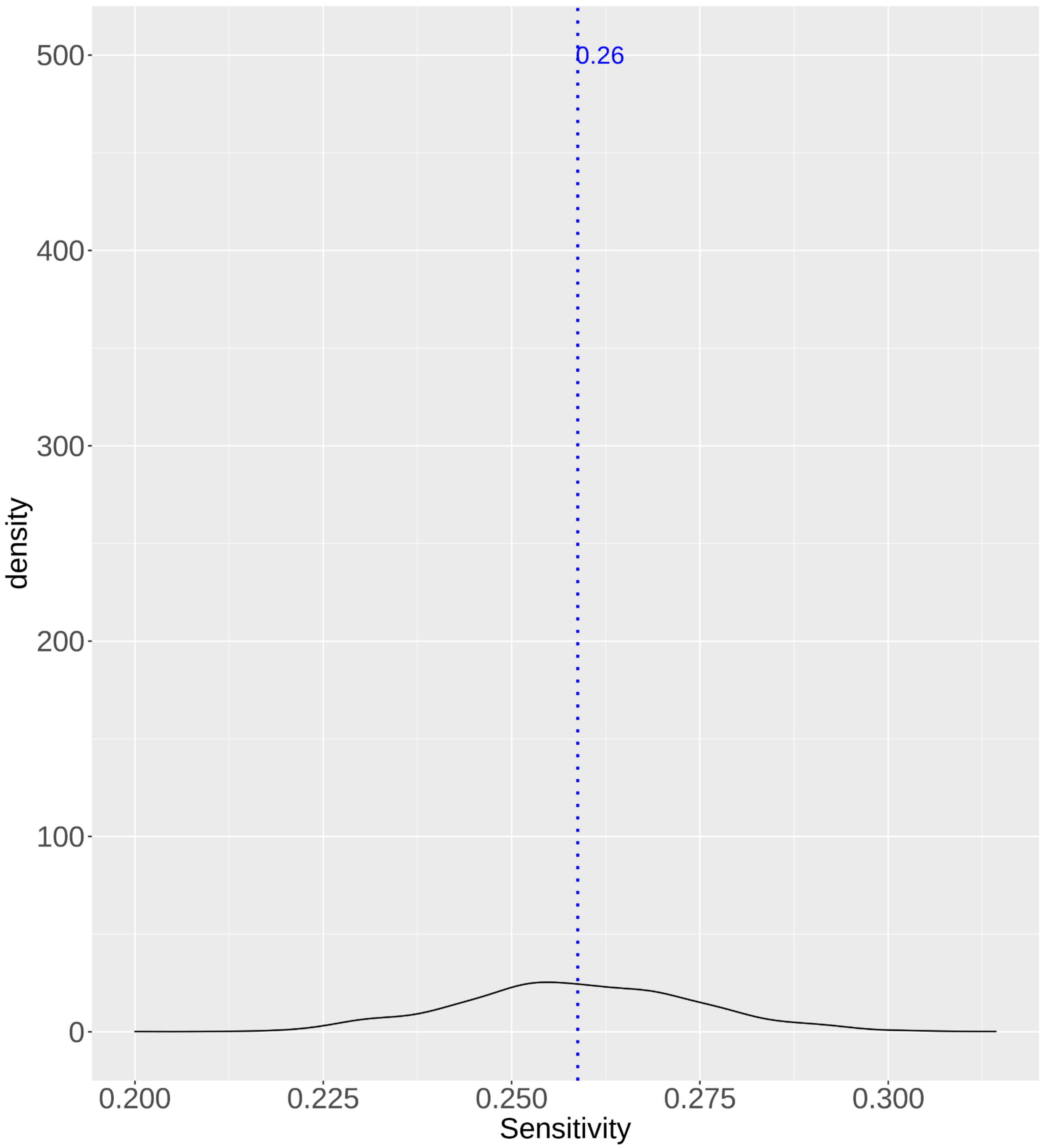


***Figure S3) Sensitivity of balanced 700 P/LP and B/LB ClinVar variants with fully conserved reference amino acids across 1,000 permutations.*** *Shown is a kernel density estimation plot of sensitivity calculated from 1000 repeated tests of a balanced set of 700 Pathogenic/Likely pathogenic (P/LP) variants that were randomly sampled and 700 Benign/Likely benign (B/LB) variants at positions that are fully conserved across the family. Average sensitivity is shown by the blue dotted line.*
